# Enhancing resolution of natural methylome reprogramming behavior in plants

**DOI:** 10.1101/252106

**Authors:** Robersy Sanchez, Xiaodong Yang, Jose R Barreras, Hardik Kundariya, Sally A. Mackenzie

## Abstract

**Background:** Natural methylome reprogramming within chromatin involves changes in local energy landscapes that are subject to thermodynamic principles. Signal detection permits the discrimination of methylation signal from dynamic background noise that is induced by thermal fluctuation. Current genome-wide methylation analysis methods do not incorporate biophysical properties of DNA, and focus largely on DNA methylation density changes, which limits resolution of natural, more subtle methylome behavior in relation to gene activity.

**Results:** We present here a novel methylome analysis procedure, Methyl-IT, based on information thermodynamics and signal detection. Methylation analysis involves a signal detection step, and the method was designed to discriminate methylation regulatory signal from background variation. Comparisons with commonly used programs and two publicly available methylome datasets, involving stages of seed development and drought stress effects, were implemented. Information divergence between methylation levels from different groups, measured in terms of Hellinger divergence, provides discrimination power between control and treatment samples. Differentially informative methylation positions (DIMPs) achieved higher sensitivity and accuracy than standard differentially methylated positions (DMPs) identified by other methods. Differentially methylated genes (DMG) that are based on DIMPs were significantly enriched in biologically meaningful networks.

**Conclusions:** Methyl-IT analysis enhanced resolution of natural methylome reprogramming behavior to reveal network-associated responses, offering resolution of gene pathway influences not attainable with previous methods.

## Background

Most chromatin changes that are associated with epigenetic behavior are reprogrammed each generation, with the apparent exception of cytosine methylation, where parental patterns can be inherited through meiosis [1]. Genome-wide methylome analysis, therefore, provides one avenue for investigation of transgenerational and developmental epigenetic behavior. Complicating such investigations in plants is the dynamic nature of DNA methylation [2, 3] and a presently incomplete understanding of its association with gene expression. In plants, cytosine methylation is generally found in three contexts, CG, CHG and CHH (H=C, A or T), with CG most prominent within gene body regions [4]. Association of CG gene body methylation with changes in gene expression remains in question. There exist ample data associating chromatin behavior with plant response to environmental changes [5], yet, affiliation of genome-wide DNA methylation with these effects, or their inheritance, remains inconclusive [6, 7].

The epigenetic landscape is modulated by thermodynamic fluctuations that influence DNA stability [8, 9] [10]. Most genome-wide methylome studies have relied predominantly on statistical approaches that ignore fundamental biophysical properties of cytosine DNA methylation, offering limited resolution of those genomic regions with highest probability of having undergone epigenetic change. Jenkinson and colleagues [11] have implemented statistical physics and information theory to the analysis of whole genome methylome data to define sample-specific energy landscapes. Our group [12, 13] proposed an information thermodynamics approach to investigate genome-wide methylation patterning based on the statistical mechanical effect of methylation on DNA molecules. The information thermodynamics-based approach is postulated to provide greater sensitivity for resolving true signal from background variation within the methylome [12]. Because gene-associated biological signal created within the dynamic methylome environment characteristic of plants may be subtle and is not free from background noise, the approach, designated Methyl-IT, includes application of signal detection theory [14–18].

A basic requirement for the application of signal detection is a probability distribution for background noise. Probability distribution, as a Weibull distribution model, can be deduced on a statistical mechanical basis for DNA methylation induced by thermal fluctuations [12]. Assuming that this background methylation variation is consistent with a Poisson process, it can be distinguished from variation associated with methylation regulatory machinery, which is non-independent for all genomic regions [12]. An information-theoretic divergence to express the background variation will follow a Weibull distribution model, provided that it is proportional to minimum energy dissipated per bit of information from methylation change.

The information thermodynamics model was previously verified with more than 150 Arabidopsis and more than 90 human methylome datasets [12]. To test application of Methyl-IT to methylome analysis, and compare resolution to approaches used in programs DSS [19], and methylpy [20], we investigated two Arabidopsis methylome datasets. For resolution of methylation signal during plant development, we used previously reported datasets from globular stage (4 days after pollination [DAP]), linear cotyledon stage (8 DAP), mature green stage (13 DAP), post-mature green stage (18 DAP), dry seed (Ws-0), and leaf [21, 22]. To assess methylation signal during stress in plants, and association of methylation with altered gene expression during stress, we investigated data from Ganguly et al. (2017), which involves mild drought stress by withholding irrigation for 9 days [23, 24]. Direct comparison of outputs by Methyl-IT with previous analyses by methylpy and DSS are presented.

## Results

### The information thermodynamics model and Methyl-IT workflow

Methylation level is generally the ratio of methylated cytosine read counts divided by the sum of methylated and unmethylated cytosine read counts for a given cytosine site. This is a descriptive variable that reflects uncertainty of methylation level at a given cytosine site. Most methylation analyses test whether or not the difference between control (CT) and treatment (TT) methylation levels (the uncertainty variation) is statistically significant. The approach measures the absolute value of the difference between methylation levels 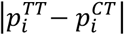 from control 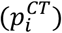 and treatment 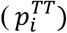 at each cytosine site. The magnitude of 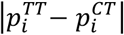 is known as total variation distance (TVD).

To improve resolution of methylation signal, we applied Hellinger divergence (*HD*), ([25], detailed description included in Methods section). Both TVD and HD are information divergences that follow asymptotic chi-square distribution [25]. However, HD converges faster and carries more information than TVD and, consequently, has higher discrimination power [26]. The improvement in discrimination power is visible in Fig. 1 By way of illustration, we used the drought stress data, where CTR designated unstressed control group and STR designated stressed group. Fig. 1a shows that treatment methylation signal on chromosome 1, expressed in terms of methylation level, was indistinguishable from control. Higher resolutions are reached with TVD and HD, with HD providing highest discrimination power.

**Fig. 1.**
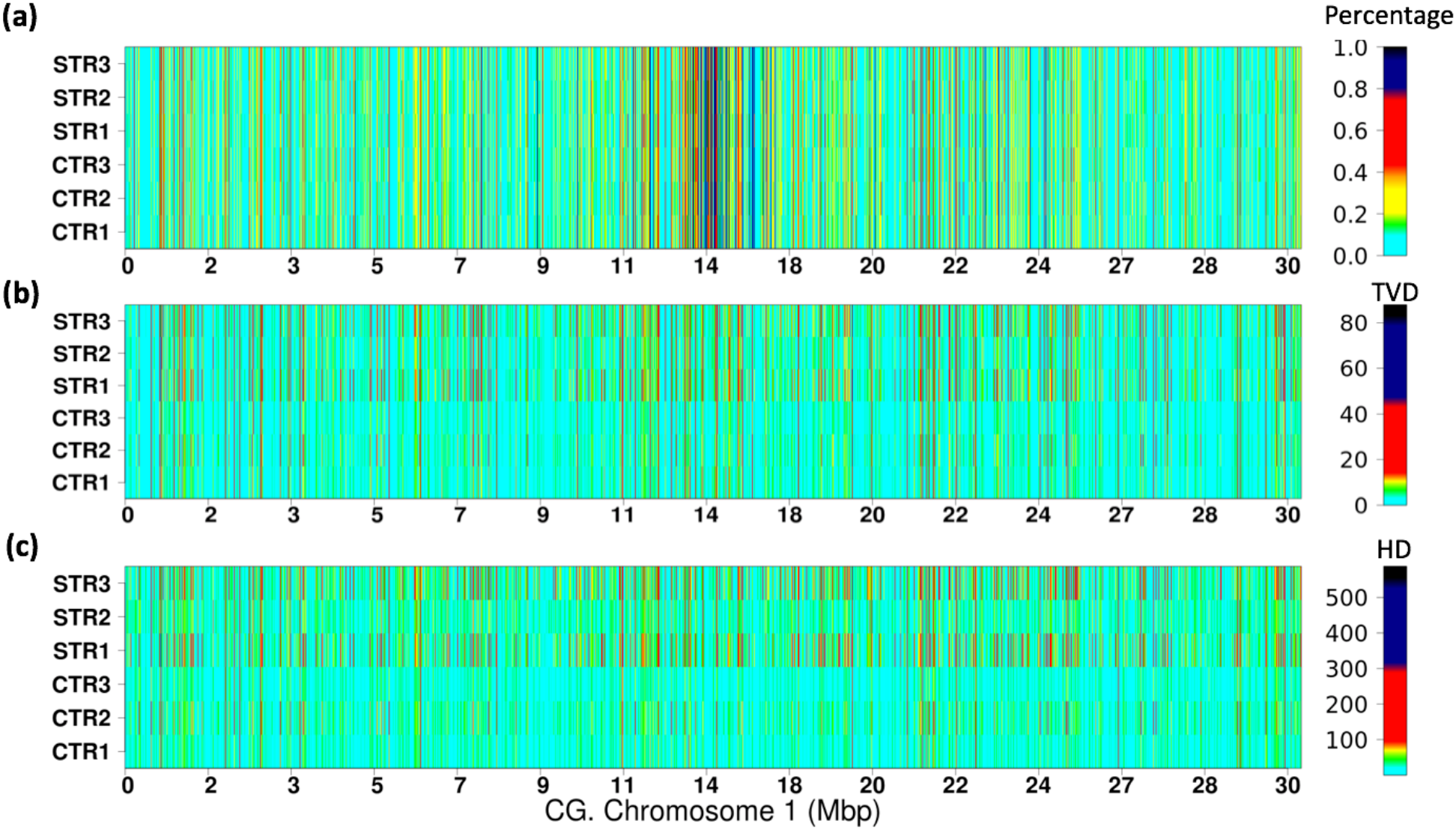
Comparison of three variables used to measure DNA cytosine methylation. The heatmap for CG methylation distribution represented by **(a)** methylation level (percentage), **(b)** total variation distance (TVD), and **(c)** Hellinger divergence (HD) on chromosome 1 for the drought stress experimental data are shown. Chromosomes were split into 2-kb non-overlapping windows (regions). The mean of methylation levels for each region *i* was estimated as: 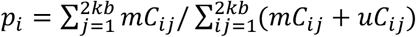 while 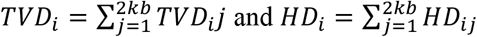.

Ganguly et al. reported individual variation and pre-existing methylation differences in the drought stress materials [24], which is reflected by HD in Fig. 1c. The improvement in resolution attributed to HD derives from the fact that TVD takes into account only one dimension of the methylation change, while HD is estimated in bi-dimensional space (*p_i_*, 1 – *p_i_*), where the goodness-of-fit test to detect differences is performed.

Genome-wide Hellinger divergence for background methylation variation can be modeled by a Weibull distribution [12]. On the other hand, biologically meaningful methylation changes result in an increment of Hellinger divergence distinguishable in the signal detection step (Fig. 2). For a given level of significance α (Type I error probability, eg. α = 0.05), cytosine positions with *H*_*α*=0.05_ can be selected as sites carrying potential biological signal (shown as the blue shade region under the curve in Fig. 2). True signal is detected based on optimal cutpoint [27], which can be estimated by area under the curve (AUC) from a receiver operating characteristic (ROC) built from logistic regression with potential signals from control and treatment. The AUC is the probability to distinguish biological regulatory signal naturally generated in the control from that induced by the treatment. Cytosine sites carrying methylation signal are designated *differentially informative methylation positions* (DIMPs). The probability that a DIMP is not induced by the treatment is designated probability of false alarm (*P_FA_*, false positive, Fig. 2). As suggested in Fig. 2, we define DIMPs as cytosine positions with high probability to carry signal created in response to treatment.

**Fig. 2.**
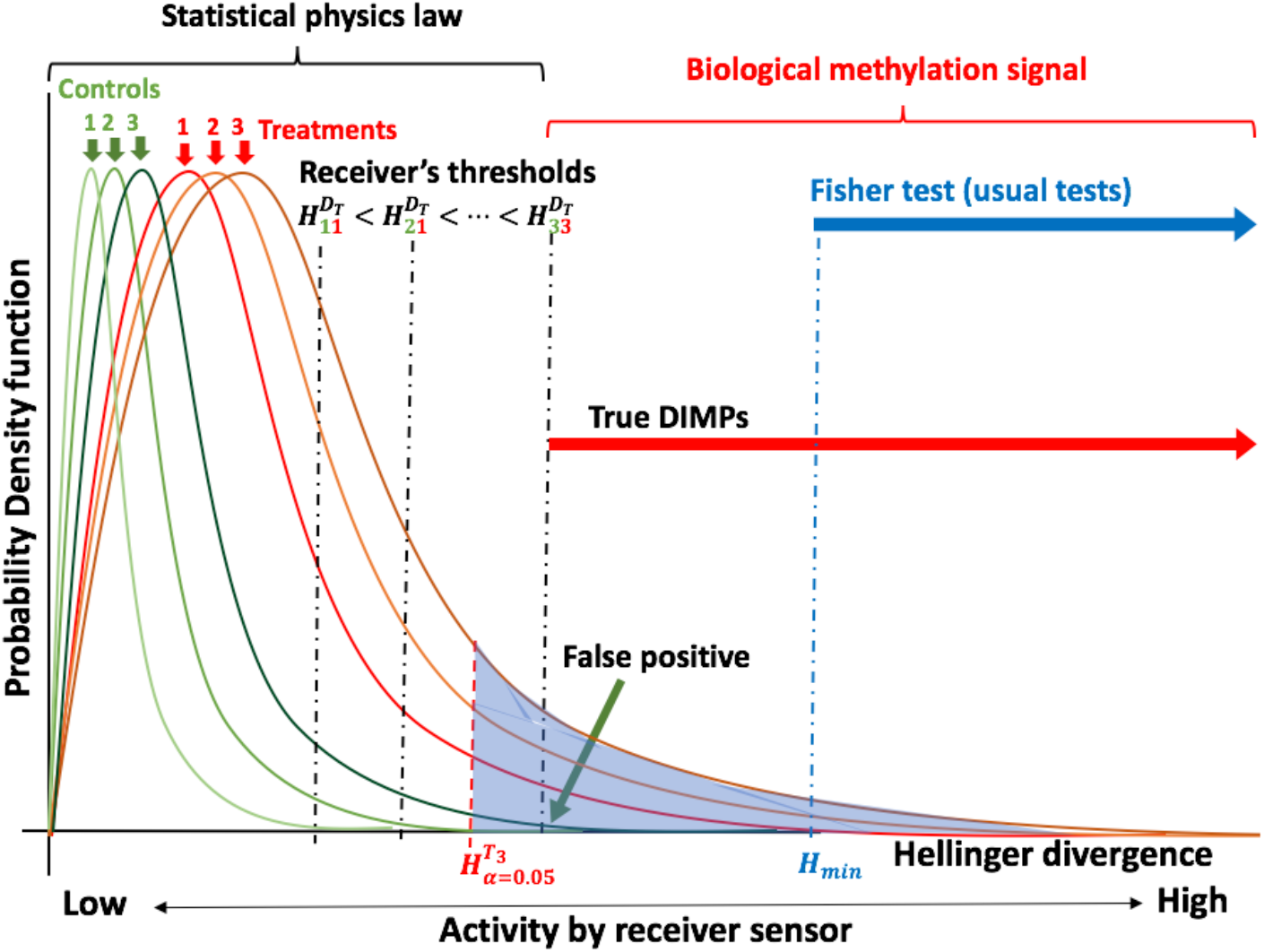
Schematic of the theoretical principle underlying Methyl-IT. Methyl-IT is designed to identify a statistically significant cutoff between thermal system noise (conforming to laws of statistical physics) and treatment signal (biological methylation signal), based on Hellinger divergence (H), to identify “true” differentially informative methylation positions (DIMPs). Empirical comparisons allow the placement of Fisher’s exact test for discrimination of DMPs.

Estimation of optimal cutoff from AUC is an additional step to remove any remaining potential methylation background noise that still remains with probability α = 0.05 > 0. We define as methylation signal (DIMP) each cytosine site with Hellinger divergence values above the cutpoint (shown in Fig. 2 as 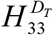). Each DIMP-associated signal may or may not be represented within a DMP derived by Fisher’s exact test (or other current tests, Fig. 2). The difference in resolution by current methods versus Methyl-IT is illustrated by positioning *H* value sensitivity for Fisher’s exact test (FET) at greater than *H_min_* for cytosine sites that are DMPs and DIMPs simultaneously (Fig. 2).

Table 1 provides a critical but non-unique example; assume there is an experiment that yields read counts with 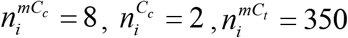, and 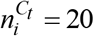, where 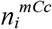 and 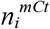 refer to methylated cytosine read counts in control and treatment, respectively, and 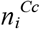 and 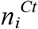 to non-methylated cytosine counts in control and treatment, respectively. In the given example, it’s clear that control and treatment have different methylation pattern, but Fisher’s exact test (including one tail test or Monte Carlo (MC) simulations with 3000 resamplings (3k)) failed to detect the difference (for significance level *α* = 0.05). Root-mean-square test (RMST) used in methylpy [20] and goodness-of-fit test based on Hellinger chi-square test (HCT, with HD as statistic) [25, 28] proved the sensitivity but still failed to detect the difference (for α= 0.05). However, if the hypothetical methylation changes were to occur in the drought stress experiment, then Weibull distribution modeling in Methyl-IT would yield *p*-values of 5.08E-04, 5.08E-04, and 3.20E-04 for each stressed plant (Table 1). Such methylation changes represent potential DIMPs. The conclusions will remain the same even for a generalized situation with 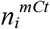 running between 80 and 350 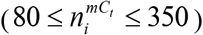. Considering that even a small genome like Arabidopsis contains millions of cytosine sites, the situation presented in Table 1 is not rare, and the difference caused by statistical tests listed in Table 1 would be significant. A flow chart integrating the main procedures of Methyl-IT and optional downstream analysis is shown in Fig. 3.

**Table 1.** Relative sensitivity differences between several statistical tests applied to identify differentially methylated cytosines. P-values for the 2×2 contingency table with read counts 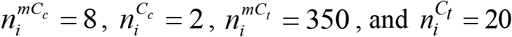

| Approach | p-value | Significance( $\alpha = 0.05$) |
| --- | --- | --- |
| FET | 0.108615 | No |
| FET one tail | 0.108615 | No |
| FET p.value MC 3k<sup>(1)</sup> | 0.1086 | No |
| RMST Boot 3k<sup>(2)</sup> | 0.051 | No |
| HT Boot 3k<sup>(3)</sup> | 0.050667 | No |
| Weibull STR1 CG<sup>(4)</sup> | 5.08E-04 | Yes |
| Weibull STR2 CG<sup>(4)</sup> | 3.20E-04 | Yes |
| Weibull STR3 CG<sup>(4)</sup> | 3.20E-04 | Yes |

<sup>1</sup>p.value simulated with Monte Carlo (MC) simulation with 3000 resamplings (3k). <sup>2</sup>Bootstrap goodness-of-fit RMST as implemented in methylpy [20]. <sup>3</sup>Bootstrap goodness-of-fit test based on Hellinger divergence estimated according to the first statistic given Theorem 1 from reference [51]. <sup>4</sup>p-value based on the Weibull distribution for memory lines (STR 1 to 3). $n_i^{mC_c}$ refers to methylated cytosine counts in control, $n_i^{C_c}$ refers to non-methylated cytosine counts in control, $n_i^{mC_t}$ refers to methylated cytosine counts in treatment and $n_i^{C_t}$ refers to non-methylated cytosine counts in treatment. The R script to compute RMST and H MC estimation is provided in GitLab: https://git.psu.edu/genomath/MethylIT

**Fig. 3.**
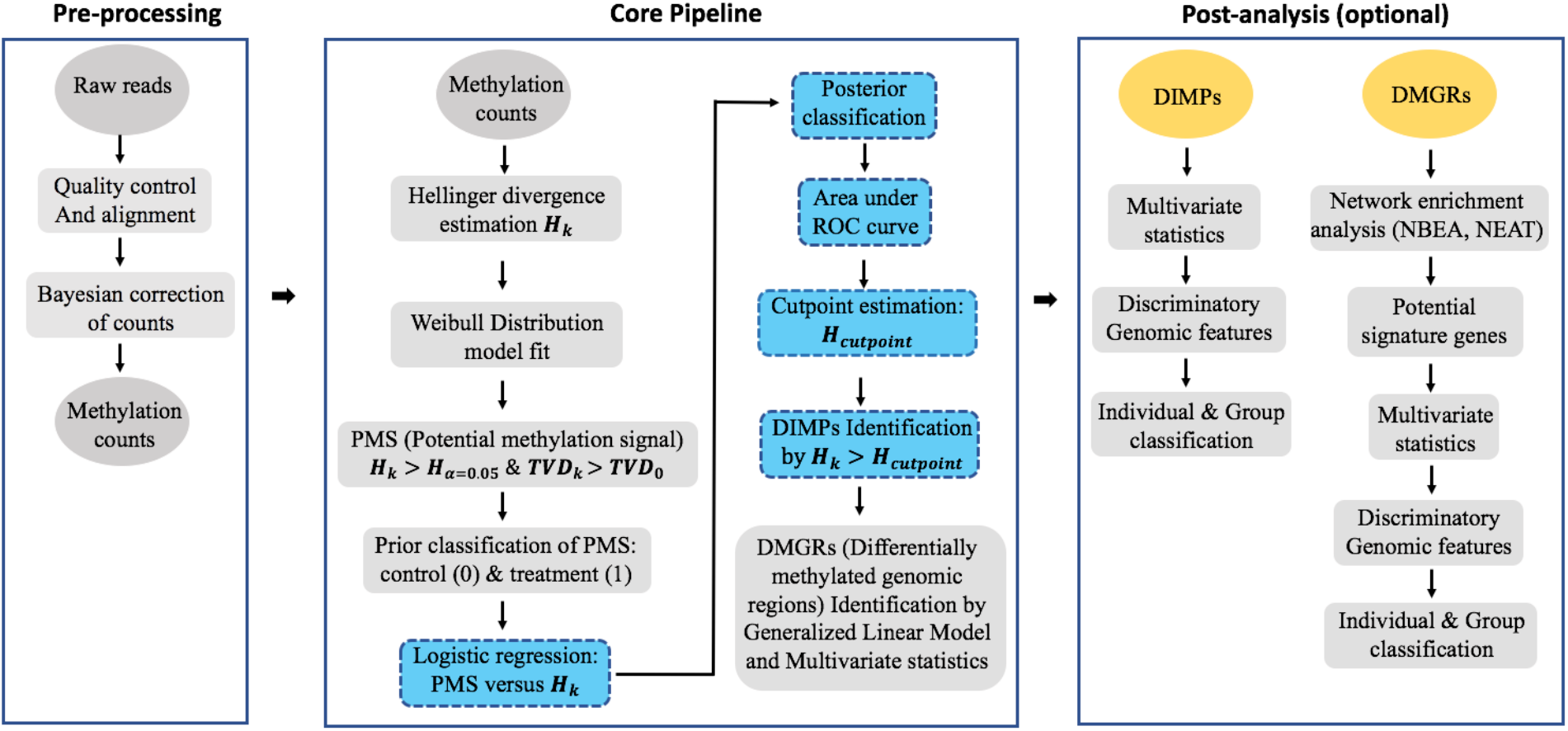
Methyl-IT processing flowchart. Ovals represent input and output data, squares represent processing steps, with signal detection processing steps highlighted in blue and DIMPs and DMGRs, as main outputs of Methyl-IT, highlighted in yellow. The generalized linear model is incorporated for group comparison of genomic regions (GRs) based on the number of DIMPs in the treatment group relative to control group. DIMPs and DMGRs can be subjected to further statistical analyses to perform network enrichment analysis and to identify potential signature genes, multivariate statistical analysis (and machine learning applications) for individual and group classifications.

### Methyl-IT sensitivity and genomic regions targeted by DIMPs

To investigate the sensitivity of Methyl-IT, we applied DIMP detection to the drought stress dataset and compared with the outputs from other methods. Fig. 4 shows a direct comparison of DIMPs to DMPs estimated with Fisher’s exact test, DMSs (differentially methylated sites) estimated with root mean square test (RMST, approach implemented in methylpy [20, 21]), and DMCs (differentially methylated cytosines) estimated with Hellinger chi-square test (HCT).

**Fig. 4.**
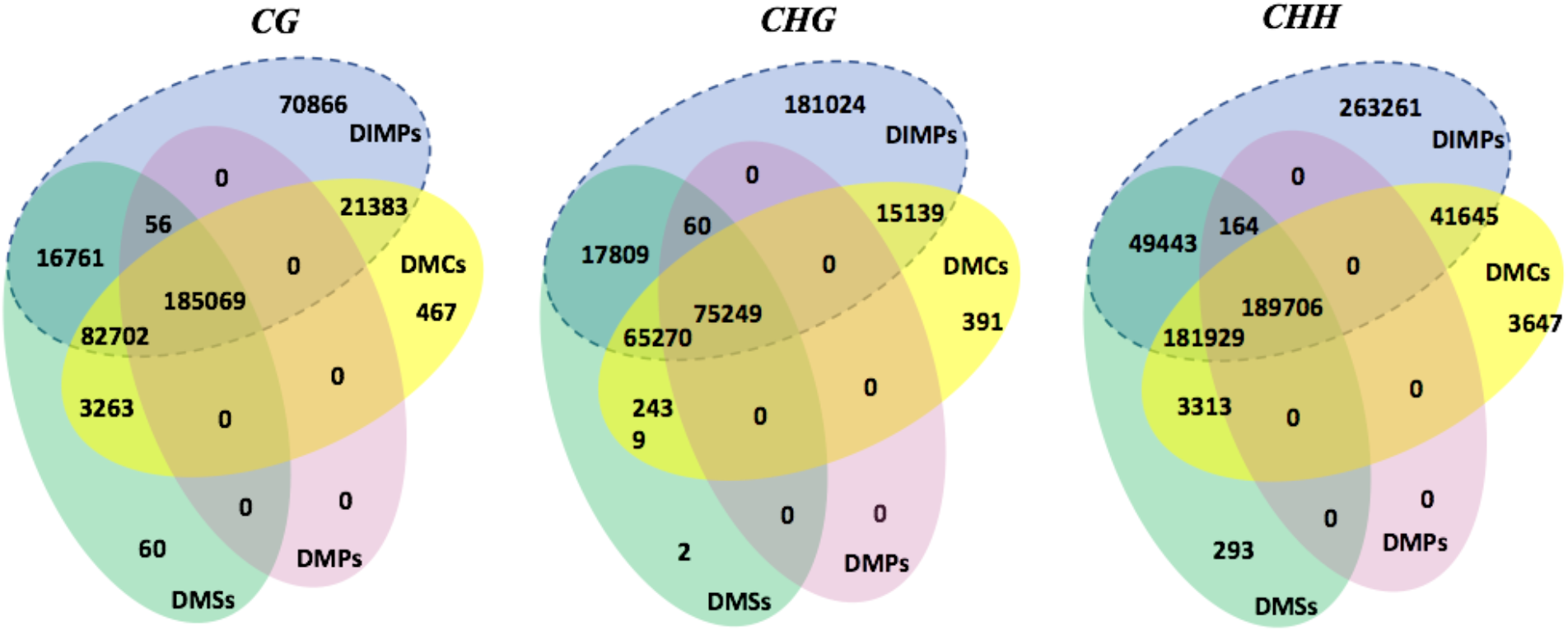
Venn diagrams of overlapping DMSs (RMST implemented in methylpy software), DMPs (obtained with Fisher Exact Test), DMCs (obtained with HCT, see methods) and DIMPs (obtained with Methyl-IT) for the drought experimental data. Only methylated cytosine positions with total variation distance (TVD) greater than 0.25 (25% of methylation level difference) are shown for the three methylation contexts. DIMPs carrying methylation signal are in the region within the dashed oval.

In all methylation contexts, 100% of DMPs (*TVD* > 0.25) found by Fisher’s exact test, 98.63% of DMSs (*TVD* > 0.25) found by RMST, and 98.45% of DMCs (*TVD* > 0.25) found by HCT are identified as DIMPs. On the other hand, DMPs only account for 30.9% of DIMPs, DMSs account for 59.8% of DIMPs, and DMCs account for 47% of DIMPs. These observations suggest a much higher sensitivity by Methyl-IT than other methods. DMS and DMC classes were relatively close, which helps validate our use of HD. Results also suggest that differences in outcome between Methyl-IT and methylpy stem from signal detection limitations rather than implementation of RMST. Application of signal detection requires knowledge of the distribution of methylation background noise, which is not a component of the methylpy procedure.

To evaluate whether DIMPs target genomic features in agreement with published reports [21–24], we assessed their distribution across the genome. Fig. 5 shows DIMP distribution pattern within three major genomic contexts (Gene regions in shades of blue, TE region in shades of red and small RNAs in shades of green). Because total cytosine number within CHH context is about 5 times higher than CG and CHG contexts, we have normalized data by presenting DIMP density (ratio of DIMP number at a given region / total cytosine context number at corresponding region) rather than absolute numbers.

**Fig. 5.**
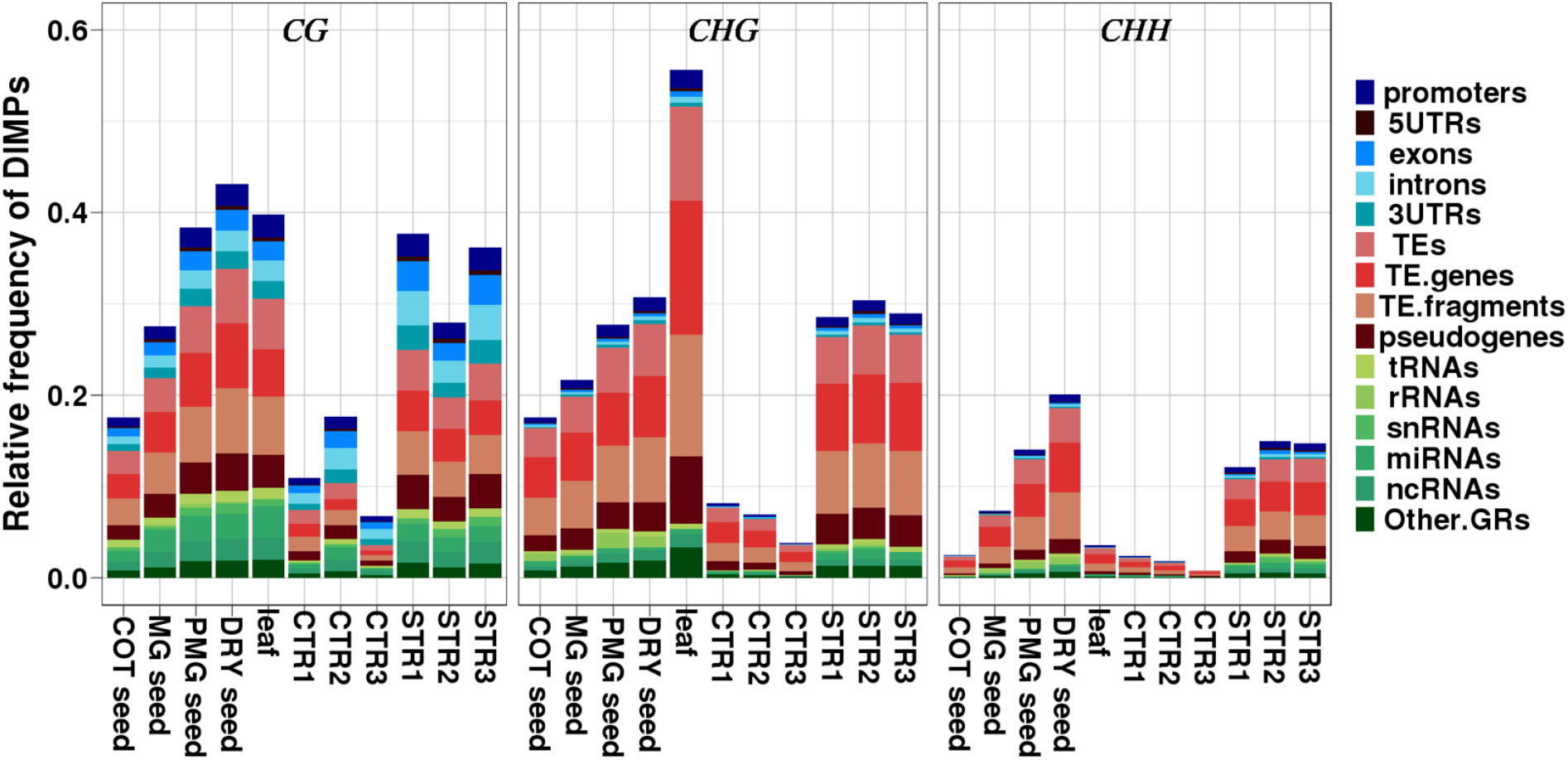
Results of signal detection with Methyl-IT for genome-wide methylome data from seed development samples from Kawakatsu et al [21] at five seed stages (GLOB, COT, MG, PMG, DRY) and leaf (globular (GLOB) stage used as control), and drought stress experiment control (CTR) and stress (STR) samples from Ganguly et al. [24]. The experimental results provide a direct, scaled comparison of methylation signal between datasets. The relative frequency of DIMPs was estimated as the number of DIMPs divided by the number of cytosine positions.

Results showed general agreement with the Kawakatsu et al. original study [21]. Strong methylation changes were identified in all three contexts during the seed development process, with DIMP signal increasing from COT to MG to PMG to dry seed, and reaching its peak in leaf tissue. CHG and CHH changes were associated predominantly with non-genic and TE regions, and CG DIMPs showed higher density within gene regions, which agreed with the DMP distribution pattern reported in the original study[21]. A surprising CHG peak was observed in leaf tissues relative to seed, which we did not pursue in detail, but may reflect a pronounced tissue-specific transition. Similar DIMP patterns were observed in the drought stress dataset relative to cytosine context, although with higher signal levels in each context.

Hierarchical clustering based on AUC criteria and built on the set of 9893 DIMP-associated genes (using *caTools* R package) permitted classification of seed developmental stages into two main phases: morphogenesis/maturation versus dormancy (Fig. 6). In this analysis, methylation signal was expressed as the sum of Hellinger divergence within genes plus 2kb upstream. Within the 9893-dimensional metric space generated by 9893 AUC-selected genes, the linear cotyledon *(COT)* and mature green (MG) stages (morphogenesis-maturation phase) grouped into a cluster quite distant from post mature green *(PMG)* and dry seed *(DRY)* stages (dormancy phase). These observations indicate a detectably greater similarity in methylome patterns between cotyledon and mature green stages, transitioning to a distinguishable state for post-mature green and dry seed. This transition may relate to the desiccation and dormancy shift that occurs within this timing [29, 30].

**Fig. 6.**
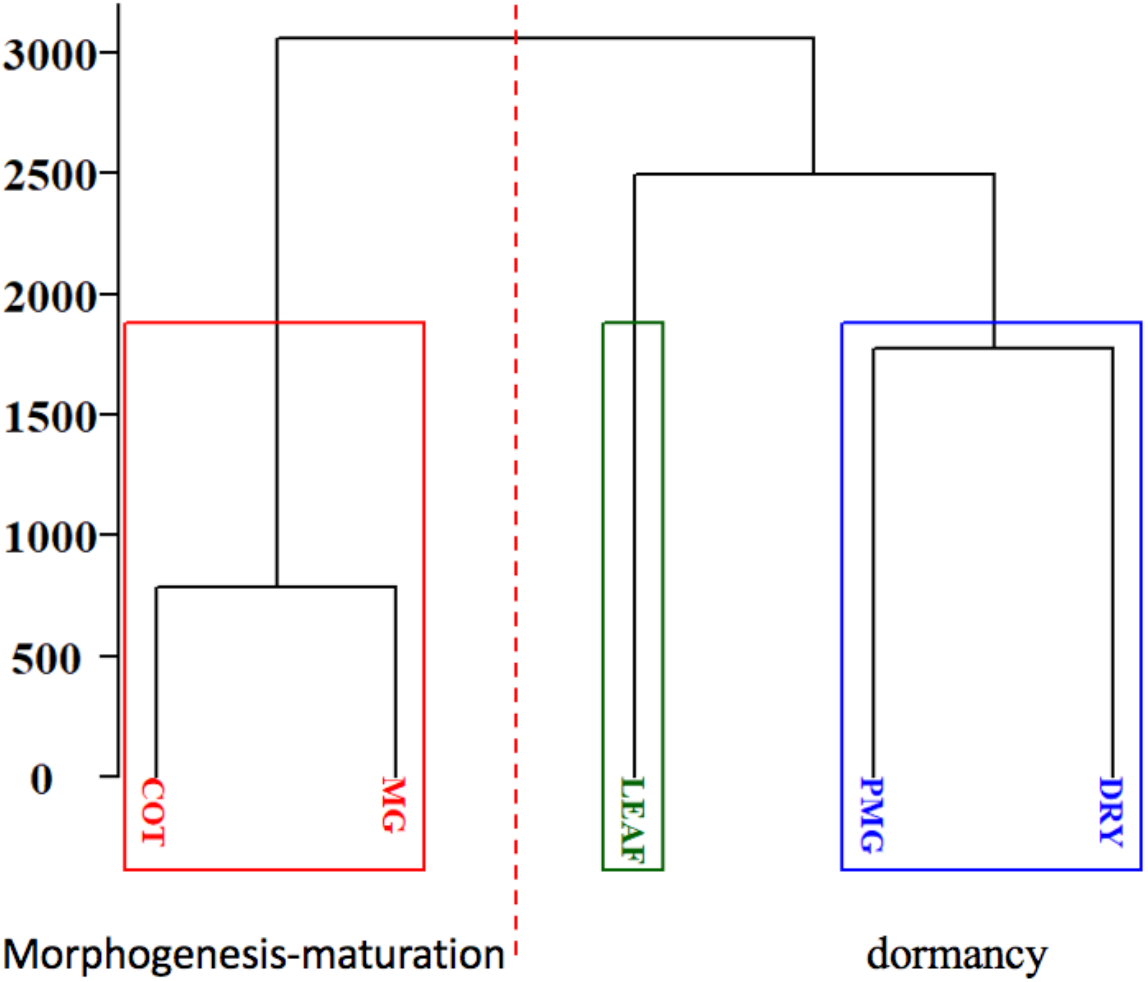
Classification of seed development stages based on identified DIMPs. A hierarchical cluster built on the set of 7006 selected DIMP-associated genes, based on AUC criteria, classified the stages into two groups: morphogenesis-maturation phase and dormancy phase.

### DIMPs can be predicted using a machine learning approach

An important test of DIMPs detected by the Methyl-IT pipeline is whether or not DIMPs identified within treatment samples can be discriminated from those in the control. To address this question, machine-learning approaches were implemented.

Each DIMP was represented as a four-dimensional vector with variables HD, TV, Weibull probability, and cytosine relative position. The classification result for simulated data and seed development data are presented in Table 2. Simulation experiments suggested that classification accuracy mainly depended on the distance separating Weibull distributions (noise plus signal) for control and treatment. Weibull model parameter values (alpha.1 and scale.1) from the first simulation for control samples (S11 to S13) were close to those estimated in the treatment group (S21 to S23), suggesting that corresponding distribution functions were close as well. Although the classifier performance to predict DIMPs could be considered acceptable (about 80% accuracy), discriminatory power to predict DIMPs from an external sample (not included to build the model) was relatively low. If probabilistic models were sufficiently distant, even a classifier trained with samples having an overall mean TVD (absolute values of methylation differences) equal to 0.13 could achieve good discrimination of DIMPs from an external sample. Importantly, a given DIMP with the same HD value in control and treatment groups could be discriminated from control group if the Weibull probability distributions from control and treatment were different.

**Table 2.** Classification of DIMPs into two classes: control (CT) and non-control (non-CT)

| Parameters from the best fitted Weibull model estimated for simulate data |  |  |  |  |  |  |  |
| --- | --- | --- | --- | --- | --- | --- | --- |
| Parameters | <sup>(1)</sup>S11 | S12 | S13 | S21 | S22 | S23 | TVD mean |
| alpha.1<sup>(2)</sup> | 0.645650 | 0.645195 | 0.645586 | 0.649747 | 0.651112 | 0.650272 | 0.03 |
| scale.1 | 0.249118 | 0.253707 | 0.253919 | 0.257382 | 0.268151 | 0.258988 |  |
| alpha.2<sup>(3)</sup> | 0.645650 | 0.645195 | 0.645586 | 0.656598 | 0.654718 | 0.652522 | 0.13 |
| scale.2 | 0.249118 | 0.253707 | 0.253919 | 0.290850 | 0.300384 | 0.290910 |  |
| Performance of classifier models built on simulated data |  |  |  |  |  |  |  |
| Classifier model | Accuracy | <sup>(4)</sup>MC. Accuracy | Sensitivity | Specificity | Pos. Pred | Neg. Pred | CT/non-CT (S23) |
| PCA-QDA.1 | 0.8088 | 0.8101 | 0.537 | 0.9763 | 0.9483 | 0.7592 | 1213/378 |
| PCA-QDA.2 | 0.9997 | 0.9998 | 1 | 0.9994 | 0.9995 | 1 | 0/2508 |
| Performance of classifier models built on CT: COT and MG, and non-CT: PMG and DRY |  |  |  |  |  |  |  |
| Methylation Context | Classif. Model | Accuracy | Sensitivity | Specificity | Pos Pred | Neg. Pred | Predictions for LEAF CT/non-CT |
| CG | Logistic | 0.9011 | 0.9984 | 0.7222 | 0.8685 | 0.996 | 18/166186 |
| CHG | Logistic | 0.7541 | 0.8842 | 0.5574 | 0.7512 | 0.7611 | 3174/205463 |
| CHH | PCA-QDA | 0.9074 | 0.9716 | 0.671 | 0.9158 | 0.865 | 69102/3906 |

<sup>(1)</sup> Simulated samples were denoted S11, S12... S23 (S11 to S13 are control, the remainder treatment). <sup>(2)</sup> 1<sup>st</sup> simulation experiment. <sup>(3)</sup> 2<sup>nd</sup> simulation experiment. <sup>(4)</sup> Accuracy mean for 500 Monte Carlos resamplings.

Classification of DIMPs was accomplished for the seed development dataset as well. Since each seed development stage comprised only one sample, groups were formed according to the hierarchical cluster presented in Fig. 6. The best classification accuracies were obtained for CG and CHH methylation contexts (Table 2). These were binary classifications, where control samples were the reference class.

Thus, probability P(x) that a new DIMP x could be observed in the control class determined its classification, and the probability that a given DIMP did not classify within the control class was 1 - P(x). A classifier model built on the groups CT: COT and MG, and non-CT: PMG and DRY (Table 2) could be applied to classify a DIMP from the leaf stage sample as non-CT. If a DIMP from the leaf stage classified as ‘CT’, this would mean that, with probability 1 – P(x) > P(x), its current methylation status for the corresponding cytosine position was not distinguishable from the status observed during early seed stages. The classifier model does not provide information for whether or not methylation status of a given cytosine position changed across the developmental stages.

### Differentially methylated genes identified by DIMPs are biologically meaningful

To investigate DIMP-based resolution of differences between seed development stages or between stressed vs non-stressed conditions, we defined differentially methylated genes (DMGs) based on group comparison for DIMP counts by applying generalized linear regression model (GLM). Genes displaying statistically significant difference in DIMP number relative to control were defined as DMGs. The DMG is defined distinctly from differentially methylated regions (DMRs), which comprise regions of high density methylation changes. In the original study of seed methylation data, enormous DMP numbers were identified in CHH context, corresponding to 23,195 DMRs that largely associated with transposable elements [21]. However, DMR association with gene regions was only scant. In the drought stress dataset, only 49 DMRs corresponding to drought stress were identified by the DSS method [24].

A total of 1068 DMGs were identified for the group comparison of morphogenesis/maturation versus dormancy phases for seed development (Additional file 1). To investigate the biological meaning of these DMGs, we conducted a network enrichment analysis test (NEAT). A statistically significant network enrichment of links between genes from the set of seed development DMGs and the set of *GO-biological process associated with seed functions* was observed (Table 3). The list of 16 networks identified includes positive and negative regulation of GA-mediated signaling, positive and negative regulation of seed germination, regulation of seed dormancy, and raffinose family oligosaccharide biosynthesis, all well-established seed processes (full gene list in Additional file 2: Table S2). GeneMANIA [31] identified interaction networks within the data, indicating that many DMGs in the seed development dataset function together (Additional file 3: Figure S1). To test the impact of different minimum cytosine coverage on Methyl-IT output, the pipeline was run without minimum coverage limit (Table 3) and with a minimum coverage of 10 reads (Additional file 4: Table S3). Results were similar with either setting.

**Table 3.** Network enrichment analysis test (NEAT) on the set of GO-biological process (BP-GO) for the differentially methylated genes in Ws-0 seed development dataset.

| BP-GO | NAB | Expected<br>NAB | Adj. *p*-<br>value |
| --- | --- | --- | --- |
| GO:0000902 cell morphogenesis | 3 | 0.2492 | 0.00280 |
| GO:0006623 protein targeting to vacuole | 4 | 0.299 | < 0.001 |
| GO:0006891 intra-Golgi vesicle-mediated transport | 4 | 0.3323 | < 0.001 |
| GO:0009723 response to ethylene | 8 | 2.9072 | 0.00873 |
| GO:0009740 gibberellic acid mediated signaling pathway | 5 | 0.9802 | 0.00375 |
| GO:0009845 seed germination | 6 | 1.3456 | 0.00301 |
| GO:0009938 negative regulation of gibberellic acid mediated signaling pathway | 4 | 0.2658 | < 0.001 |
| GO:0010162 seed dormancy process | 5 | 1.03 | 0.00434 |
| GO:0010187 negative regulation of seed germination | 3 | 0.4319 | 0.00916 |
| GO:0010325 raffinose family oligosaccharide biosynthetic process | 5 | 0.3323 | 0.00102 |
| GO:0016049 cell growth | 3 | 0.3655 | 0.00640 |
| GO:0016192 vesicle-mediated transport | 5 | 0.3987 | < 0.001 |
| GO:0016197 endosomal transport | 2 | 0.0665 | 0.00280 |
| GO:0048444 floral organ morphogenesis | 5 | 0.3323 | < 0.001 |
| GO:2000033 regulation of seed dormancy process | 3 | 0.1994 | 0.0017 |
| GO:2000377 regulation of reactive oxygen species metabolic process | 4 | 0.4153 | 0.00154 |

Only over-enriched pathways are included
NAB: observed number of (network) links from DMG list to GO term gene list
Expected NAB: expected number of links from DMG list to GO term gene list (in absence of enrichment)
Enrichment Fold: the ratio of NAB (observed number of network links) / expected nab (expected number of links)

**Table 4.**
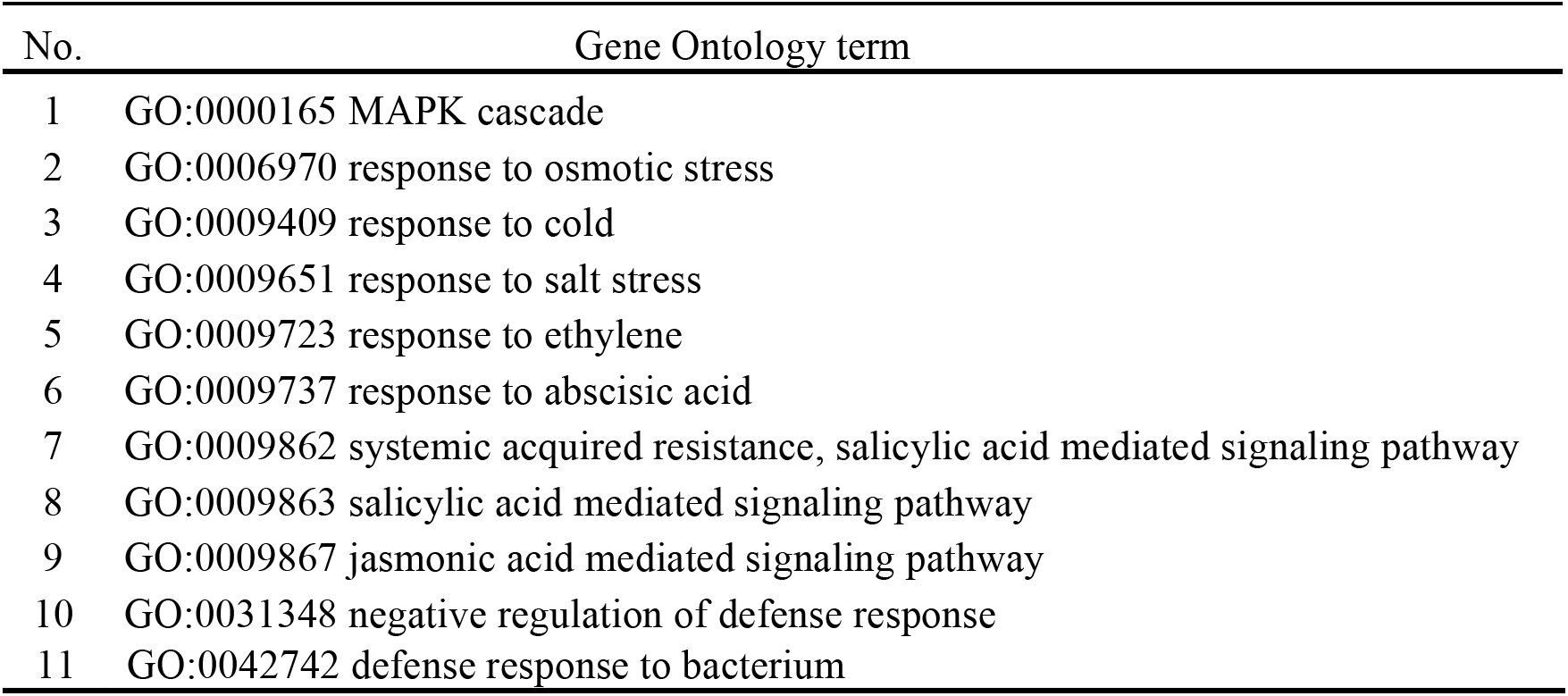
Overlapped pathways between DEGs and DMGs in drought stress data

In the drought stress experiment, analyses performed by the original authors detected 2141 CG, 1039 CHG and 718 CHH DMRs, which eventually led to identification of 49 drought stress-related DMRs [24]. A very weak relationship between methylome changes and phenotype or gene expression patterns was suggested in the original study [24]. With Methyl-IT, we identified 6669 DMGs (Additional file 5: Table S4). To investigate whether associations between identified DMGs and gene expression were evident, we compared the DMG list with the differentially expressed gene (DEG) dataset reported in the original study with 4371 genes [23]. Fig. 7a shows that the two lists shared 842 genes, accounting for 19.25% DEGs and 12.6% DMGs. Applying NEAT and Network Based Enrichment Analysis (NBEA) to DEG and DMG datasets, we identified 73 significantly enriched DEG and 23 DMG networks. Among them, 11 were shared and all were related to plant stress response mechanisms. Fig. 8 shows four examples within the 11 networks, with MAPK cascade (GO:0000165), response to osmotic stress (GO:0006970), response to salt stress (GO:0009651), and response to abscisic acid (GO:0009737). Each gene shown carried significant DIMP signal (Additional file 5: Table S4, Additional file 6: Table S5), suggesting that a systematic methylation repatterning had occurred within these networks. At an individual gene level, numerous genes showed both significant gene expression and methylation changes associated with drought stress response. For example, *ABA INSENSITIVE 1 (ABI1,* AT4G26080) encodes a protein involved in abscisic acid signal transduction that negatively regulates ABA promotion of stomatal closure [32]. The locus carries 5 DIMPs on average in the three drought stressed plants, and is up-regulated 3.36 fold. *ABRE BINDING FACTOR 4 (ABF4,* AT3G19290), encodes a bZIP transcription factor with specificity for abscisic acid-responsive elements (ABRE), and mediates ABA-dependent stress responses, acting through the SnRK2 pathway [33]. This gene has an average of 8.7 DIMPs and 2.6-fold up-regulation. *ABSCISIC ACID RESPONSIVE ELEMENTS-BINDING FACTOR 3 (ABF3,* AT4G34000) encodes an ABA-responsive element-binding protein with similarity to transcription factors expressed in response to stress and abscisic acid [34]. In our study, this gene displays 11 DIMPs and is up-regulated 11 fold. *CIRCADIAN CLOCK ASSOCIATED 1 (CCA1,* AT2G46830) encodes a transcriptional repressor that performs overlapping functions with *LHY* in a regulatory feedback loop that is closely associated with the circadian oscillator of Arabidopsis [35]. This gene shows an average of 7.7 DIMPs and is up-regulated 38.6 fold. Taken together, these data provide enticing indication that differential gene methylation is subtle, goes undetected by common methodologies, and identifies gene networks that are compelling candidates for more detailed subsequent investigation.

**Fig. 7.**
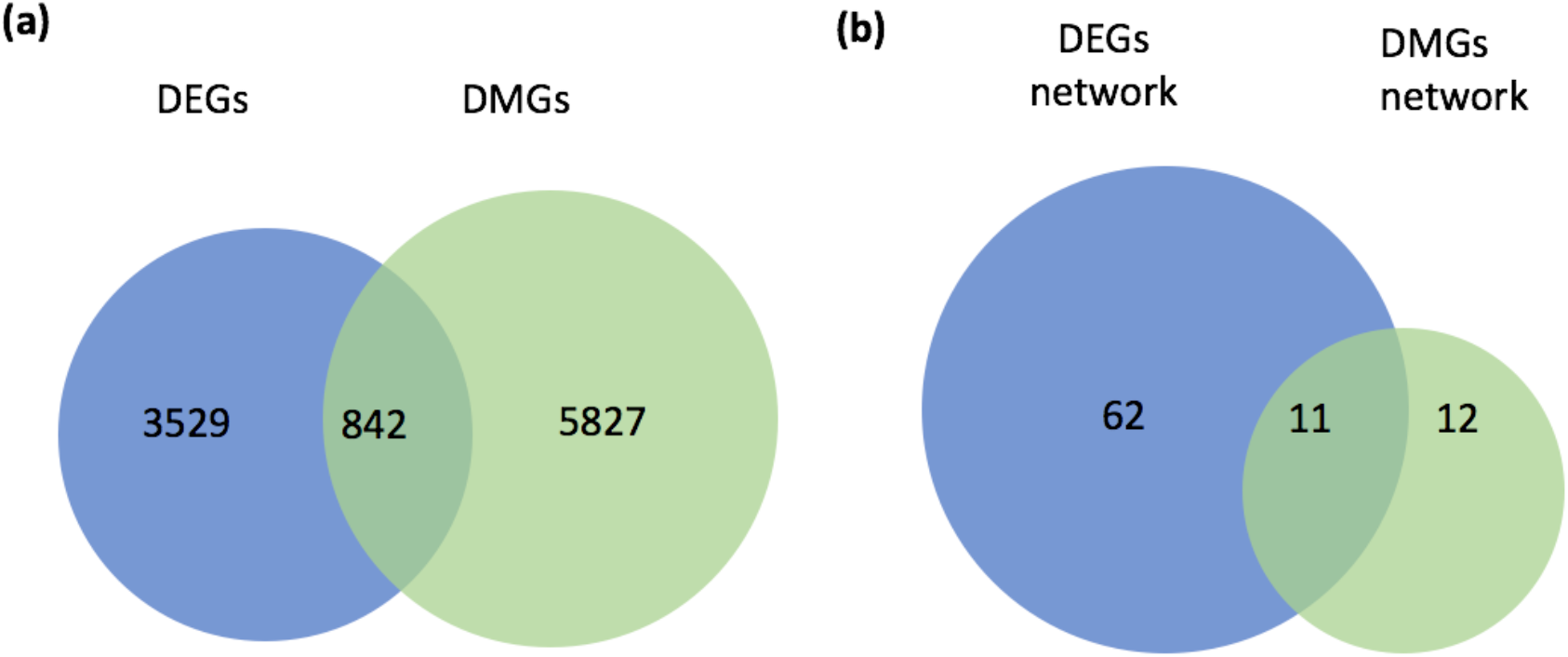
Differential expressed genes(DEGs) vs differential methylated genes(DMGs) in unstressed plants vs drought stressed plants comparison. **(a)** 4371 DEGs were identified by [24] and 6669 DMGs were identified by Methyl-IT. **(b)** 73 and 23 significantly enriched networks were identified from 4371 DEGs and 6669 DMGs, respectively. NEAT and NBEA analysis were used to identify enriched network (see Method).

**Fig. 8.**
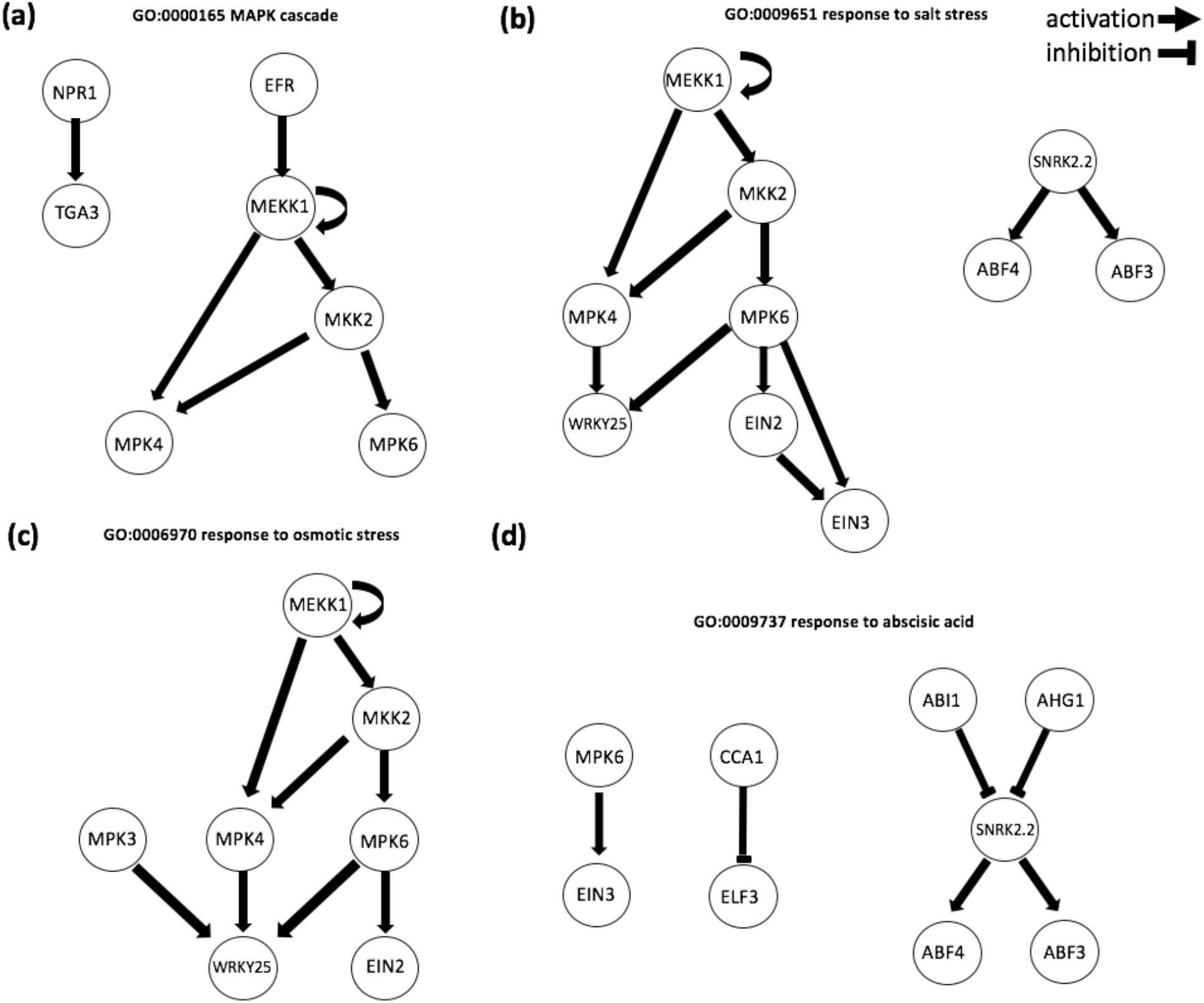
Examples of enriched networks identified by NBEA using DMGs. Genes involved in **(a)** MAPK cascade, **(b)** response to salt stress, **(c)** response to osmotic stress, and **(d)** response to abscisic acid are in circles. Network graphs were generated by EnrichmentBrowser R package in R. Details of DIMPs number for each gene could be found in additional file 6.

## Discussion

Methyl-IT draws from the perspective that DNA methylation functions to stabilize DNA [8, 36, 37] and, as such, may exist in “activated-signal” versus “maintenance” states with regard to bioenergetics. The theoretical premise underlying our approach, and based on Landauer’s principle, is detailed elsewhere [12, 13], while the present study compares resolution of this methodology to current methods for analysis of whole-genome methylation datasets. To date, there has not been a statistical biophysics model to simulate background methylome variation. Consequently, comparisons with other methylation analysis procedures presented here were limited to published experimental datasets.

Methyl-IT permits methylation analysis as a signal detection problem. The model predicts that most methylation changes detected, at least in Arabidopsis, represent methylation “background noise” with respect to methylation regulatory signal, explainable within a statistical physical probability distribution. Implicit in our approach is that DIMPs can be detected in the control sample as well. These DIMPs are located within the region of false alarm in Fig. 1, and correspond to natural methylation signal not induced by treatment. Thus, using the Methyl-IT procedure, methylation signal is not only distinguished from background noise, but can be used to discern natural signal from that induced by treatment.

Whereas methods underlying RMST (methylpy approach) and DSS provide essential information about methylation density, context and positional changes on a genome-wide scale, Methyl-IT provides resolution of subtle methylation repatterning signals distinct from background fluctuation. Data derived from analysis with FET, RMST, HCT or DSS alone could lead to an assumption that gene body methylation plays little or no role in gene expression, or that transposable elements are the primary target of methylation repatterning. Yet ample data suggest that this picture is incomplete [38]. Methyl-IT results show that these conclusions more likely reflect inadequate resolution of the methylome system. GLM analysis applied to the identification of DMR-associated genes by methylpy [21] and DSS indicates that DMRs (or DMR associated genes) do not provide sufficient resolution to link them with gene expression.

Signal detected by Methyl-IT may reflect gene-associated methylation changes that occur in response to local changes in gene transcriptional activity. Pathway-associated methylome changes detected in seed development data suggest participation of methylation in gene expression stage transitions, particularly prominent between mature green and post-mature green stages. Likewise, coincident patterns between methylome-associated gene networks and gene expression networks during drought stress appear to be strongly non-random.

Methyl-IT analysis of various stages in seed development and germination showed evidence of methylation changes. Previous methylpy output [21] defined predominant changes in non-CG methylation residing within TE-rich regions of the genome, whereas Methyl-IT data resolved statistically significant methylation signal within gene regions. With the complementary resolution provided by Methyl-IT, it becomes possible to investigate the nature of chromatin response within identified genes in greater detail during the various stages of a seed’s development. Several of the identified DMGs in this study involved genes that interact within known seed-associated pathways.

A limitation to currently existing methylome data analysis platforms is that most require fairly advanced coding skills and statistics knowledge, rendering them less directly accessible to most biologists. Methyl-IT has been designed to be highly user friendly, accessible to any biologist with basic R knowledge.

## Conclusions

Methyl-IT is an alternative and complementary approach to plant methylome analysis that discriminates DNA methylation signal from background and enhances resolution. Analysis of publicly available methylome datasets showed enhanced signal during seed development and germination or during drought stress within genes belonging to related pathways, providing new evidence that DNA methylation changes occur within gene networks. Whereas, previous methylome analysis protocols identify changes in methylome density and landscape, predominantly non-CG, Methyl-IT reveals effects within gene space, mostly CG and CHG, for elucidation of methylome linkage to gene effects.

## Methods

### Methylome analysis

The alignment of BS-Seq sequence data from *Arabidopsis thaliana* was carried out with Bismark 0.15.0 [39]. BS-Seq sequence data from tomato experiment were aligned using ERNE 2.1.1 [40]. The basic theoretical aspects of methylation analysis applied in the current work are based on previous published results [12]. Details on Methyl-IT steps are provided in the next sections.

#### Methylation level estimation

In Methyl-IT pipeline, it is up to the user whether to estimate methylation levels at each cytosine position following a Bayesian approach or not. In a Bayesian framework assuming uniform priors, the methylation level *P_i_* can be defined as: 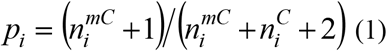, where 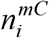 and 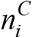 represent the numbers of methylated and non-methylated read counts observed at the genomic coordinate *i*, respectively. We estimate the shape parameters *α* and *β* from the beta distribution 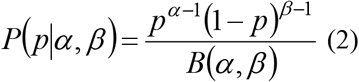 minimizing the difference between the empirical and theoretical cumulative distribution functions (ECDF and CDF, respectively), where *B*(*α, β*) is the beta function with shape parameters *α* and *β*. Since the beta distribution is a prior conjugate of binomial distribution, we consider the *p* parameter (methylation level *P_i_*) in the binomial distribution as randomly drawn from a beta distribution. The hyper-parameters *α* and *β* are interpreted as pseudo counts. Then, the mean 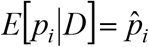 of methylation levels *p_i_*, given the data *D*, is expressed by 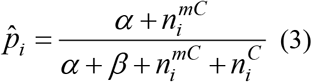. The methylation levels at the cytosine with genomic coordinate *i* are estimated according to this equation. If the Bayesian framework is not selected, then methylation levels are estimated as: 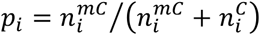.

#### Hellinger and Total Variation divergences of the methylation levels

To evaluate the methylation differences between individuals from control and treatment we introduce a metric in the bidimensional space of methylation levels: *P_i_* = (*p_i_*, 1 – *p_i_*). Vectors *P_i_* provide a measurement of the uncertainty of methylation levels at position *i*. However, we do not perform a direct comparison between the uncertainty of methylation levels from each group of individuals, control 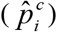 and treatment 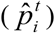, but the uncertainty variation with respect to the same individual reference 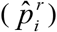 on the mentioned metric space. The reason to measure the uncertainty variation with respect to the same reference resides in that even sibling individuals follow an independent ontogenetic development. This a consequence of the “omnipresent” action of the second law of thermodynamics in living organisms, at molecular level manifested throughout the actions of Brownian motion and thermal fluctuations on DNA molecules.

The difference between methylation levels from reference and treatment (control) experiments is expressed in terms of information divergences of their corresponding methylation levels,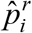 and 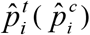, respectively. The reference sample(s) can be additional experiment(s) fixed at specific conditions, or a virtual sample created by pooling methylation data from a set of control experiments, e.g. wild type individual or group.

If the read counts 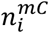 and 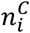 are provided and taken into account, then the Hellinger divergence between the methylation levels from reference and treatment experiments is defined as:

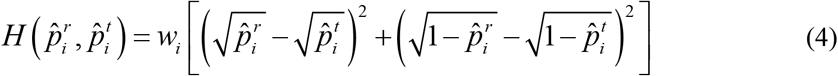

Where 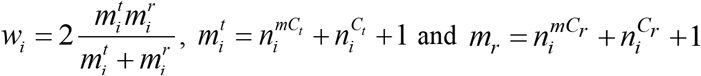. Otherwise, Hellinger divergence between the methylation levels from reference and treatment experiments is defined as:

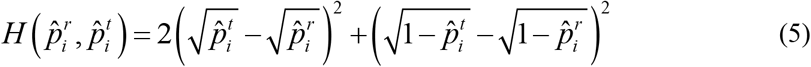

The total variation of the methylation levels 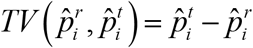 (6) indicates the direction of the methylation change in the treatment, hypo-methylated *TV* < 0 or hyper-methylated *TV* > 0. *TV* is linked to a basic information divergence, the total variation distance, defined as: 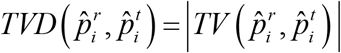 (7) [41]. Distance 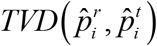 and Hellinger divergence (as given in Eq. 4) hold the inequality: 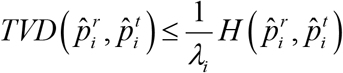 (8), where *λ_i_* = *w_i_*/2, which is a direct consequence of the Cauchy-Schwarz inequality. Under the null hypothesis of non-difference between distributions 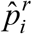 and 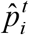, Eq. 4 asymptotically has a chi-square distribution with one degree of freedom, which set the basis for a Hellinger chi-square test (HCT). The term *w_i_* introduces a useful correction for the Hellinger divergence, since the estimation of 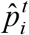 and 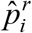 are based on counts (see Table 1).

In Methyl-IT pipeline, the statistics mean, median, or sum of the read counts at each cytosine site of some control samples can be used to create a virtual reference sample. It is up to the user whether to apply the ‘row sum’, ‘row mean’ or ‘row median’ of methylated and unmethylated read counts at each cytosine site across individuals.

#### Non-linear fit of Weibull distribution

The cumulative distribution functions (CDF) for 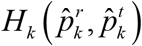 can be approached by a Weibull distribution 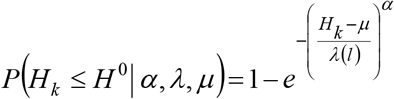 (9) [12]. Parameter 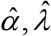 and 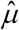 were estimated by nonlinear regression analysis of the ECDF 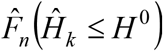 versus 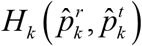 [12]. The ECDF of the variable 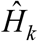 is defined as:

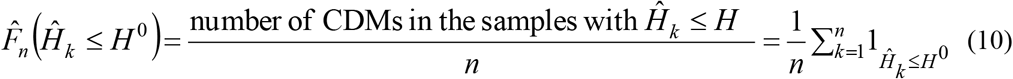

where 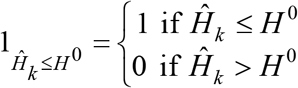 is the indicator function. Function 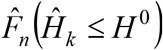 is easily computed (for example, by using function “*ecdf*’ of the statistical computing program “R”[42]).

#### A statistical mechanics-based definition for a potential/putative methylation signal (PMS)

Most methylation changes occurring within cells are likely induced by thermal fluctuations to ensure thermal stability of the DNA molecule, conforming to laws of statistical mechanics [12]. These changes do not constitute biological signals, but methylation background noise induced by thermal fluctuations, and must be discriminated from changes induced by the treatment. Let 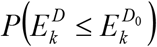 be the probability that energy 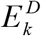, dissipated to create an observed divergence *D* between the methylation levels from two different samples at a given genomic position *k*, can be lesser than or equal to the amount of energy 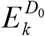. Then, a single genomic position *k* shall be called a PMS at a level of significance *α* if, and only if, the probability 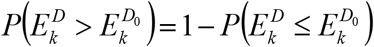 to observe a methylation change with energy dissipation higher than 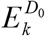 is lesser than *α*. The probability 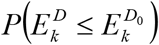 can be given by a member of the generalized gamma distribution family and, in most cases, experimental data can be fixed by the Weibull distribution [12]. Based on this dynamic nature of methylation, one cannot expect a genome-wide relationship between methylation and gene expression. A practical definition of PMS based on Hellinger divergence derives provided that *H_k_* is proportional to 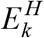 and using the estimated Weibull CDF for *H_k_* given by Eq. 8. That is, a single genomic position *k* shall be called a PMS at a level of significance *α* if, and only if, the probability 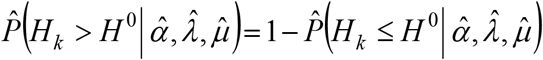 to observe a methylation change with Hellinger divergence higher than *H_k_* is lesser than *α*.

The PMSs reflect cytosine methylation positions that undergo changes without discerning whether they represent biological signal created by the methylation regulatory machinery. The application of signal detection theory is required for robust discrimination of biological signal from physical noise-induced thermal fluctuations, permitting a high signal-to-noise ratio [18].

#### Robust detection of differentially informative methylated positions (DIMPs)

Application of signal detection theory is required to reach a high signal-to-noise ratio [43, 44]. To enhance DIMP detection, the set of PMSs is reduced to the subset of cytosines with 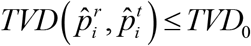, where *TVD*_0_ is a minimal total variation distance defined by the user, preferably *TVD*_0_ > 0.1. If we are interested not only in DIMPs but also in the full spectrum of biological signals, this constraint is not required. Once potential DIMPs are estimated in the treatment and in the control samples, a logistic regression analysis is performed with the prior binary classification of DIMPs, i.e., in terms of PMSs (from treatment versus control), and a receiver operating curve (ROC) is built to estimate the cutpoint of the Hellinger divergence at which an observed methylation level represents a true DIMP. There are several criteria to estimate the optimal cutpoint, many of which are implemented in the R package *OptimalCutpoints* [27]. The optimal cutpoint used in Methyl-IT corresponds to the *H* value that maximizes Sensitivity and Specificity simultaneously [45, 46]. These analyses were performed with the R package *Epi* [47].

Once all pairwise comparisons are done, a final decision of whether a DFMP is a DIMP is taken based on the highest cutpoint detected in the ROC analyses (Fig. 1). That is, the decision is taken based on the cutpoint estimated in the ROC analysis for the control sample with the closest distribution to treatment samples. The position of the cutpoint will determine a final posterior classification for which we would estimate the number of true positive, true negatives, false positives and false negatives. For each cutpoint we would estimate, the accuracy and the risk of our predictions. We may wish to use different cutpoints for different situations. For example, if our goal is the early detection of a terminal disease and high values of the target variable indicates that a patient carries the disease, then to save lives we would prefer the lowest meaningful cutpoint reducing the rate of false negative.

#### DIMP simulation and machine learning classifier

Methyl-IT pipeline was applied to seven random generated individual samples, each on with 2×10^5^ simulated cytosine positions with their corresponding methylation levels. A reference individual sample was generated with parameters *α* = 1.54 and *β* = 2 with mean of methylation levels 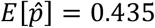 and variance 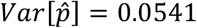. Two simulation experiments were performed. For the first simulation, total variations values for three control samples (S11 to S13) were generated using normal distribution with means (standard deviation): 0.297 (0.31), 0.297 (0.32), and 0.295 (0.34) and for three treatment (S21 to S23) individual with means (standard deviation): 0.44 (0.3), 0.45 (0.33), and 0.43 (34). The overall mean of all the pairwise differences of methylation levels between control and treatment sample is 0.03.

TV treatment means were increase in the second simulation with values: 0.54, 0.55, and 0.53. The overall mean of all the pairwise differences of methylation levels between control and treatment sample is 0.13. DIMPs were estimated according to Methyl-IT pipeline and a classifier model was inferred with the three control samples and the first two treatment samples to classify DIMPs into two classes: control (CT) and treatment (non-CT or ‘TT’). Each cytosine site is represented as a four dimensional vector with variables: HD, TV, Weibull probability, and cytosine relative position estimated as (*x* – *x_min_*)/(*x_max_* – *x*), where *x_min_* and *x_max_* are the maximum and minimum positions for the corresponding chromosome.

The set of four dimensional vectors integrated by control and treatment was randomly split into two subsets: training (60%, used to train the model) and test (40%, used to evaluate the classifier). The classification performance was evaluated with Monte Carlo resampling and the classifier model was applied to predict DIMPs from the third treatment sample not included in the construction of the classifier model. In the case of Monte Carlo resampling, a new random split of the samples is performed for each resampling.

Currently, there are seven classifiers available to use with Methyl-IT: logistic regression model (LRM), linear discriminant analysis (LDA), quadratic discriminant analysis (QDA), support vector machine (SVM), PCA-LRM using the principal component (PCA) as predictor variables in LMR, PCA-LDA and PCA-QDA.

#### Estimation of differentially methylated genes (DMGs) using Methyl-IT

Our degree of confidence in whether DIMP counts in both control and treatment represent true biological signal was set out in the signal detection step. To estimate DMGs, we followed similar steps to those proposed in Bioconductor R package DESeq2 [48], but the test looks for statistical difference between the groups based on gene body DIMP counts rather than read counts. The regression analysis of the generalized linear model (GLMs) with logarithmic link was applied to test the difference between group counts. The fitting algorithmic approaches provided by *glm* and *glm.nb* functions from the R packages *stat* and MASS were used for Poisson (PR), Quasi-Poisson (QPR) and Negative Binomial (NBR) linear regression analyses, respectively.

Likewise for DESeq2 we used the linear regression model 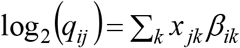, with design matrix elements *x_jk_*, coefficients *ß_ik_*, and mean *μ_kj_* = *S_j_q_kj_*, where *S_j_* normalization constants are considered constant within a group. Only two groups were compared at a time. The design matrix elements indicate whether a sample *j* is treated or not, and the GLM fit returns coefficients indicating the overall methylation strength at the gene and the logarithm base 2 of the fold change (log2FC) between treatment and control [48]. In particular, in the case of NBR, the inverse of the variance was used as prior weight (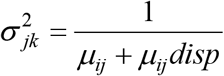, where *disp* is data dispersion computed by the *estimateDispersions* function from DESeq2 R package).

To test difference between group counts we applied the fitting algorithmic approaches: PR and PQR if 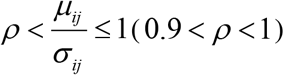, NBR and NBR with ‘*prior weights*’. Next, best model based on Akaike information criteria (AIC). The Wald test for significance of the independent variable coefficient indicates whether or not the treatment effect is significant, while the coefficient sign (*log2FC*) will indicate the direction of such an effect.

#### Bootstrap goodness-of-fit test for 2310 cl:432 contingency tables

The goodness-of-fit RMST 2311 cl:292 contingency tables as implemented in methylpy [20] for the estimation of DMSs (based on the root-mean-square (RMS) statistics) is explained in Perkins et al. in reference [49](a complemental description is found at arXiv: 1108.4126v2). The bootstrap heuristic to perform the test is given in reference [50]. An analogous bootstrap goodness-of-fit test based on Hellinger divergence was also applied to estimate DMCs. In this case, Hellinger divergence estimated according to the first statistic given in Theorem 1 from reference [51].

#### Network enrichment analysis

Network based enrichment analysis (NBEA) was applied using the EnrichmentBrowser R package [52, 53] and the Network Enrichment Analysis Test (NEAT) was performed by using the R package “neat” version 1.1.1[53].

## Abbreviations

AUC: Area under the receiver operating characteristic curve
CDM: Cytosine DNA methylation
DAGs: DMR associated genes
DEG: Differentially expressed gene
DIMPs: Differentially informative methylated positions
DMGs: Differentially methylated genes
DMPs: Differentially methylated positions
DMRs: differentially methylated regions
DSS: Dispersion Shrinkage for Sequencing
FET: Fisher’s exact test
GLM: generalized linear regression model
HD: Hellinger divergence
HCT: Hellinger chi-square test. Goodness-of-fit test based on Hellinger divergence
NEAT: Network Enrichment Analysis Test
NBEA: Network based enrichment analysis
RMST: Root-mean-square test
ROC: Receiver operating characteristic curve
SD: Signal detection
TVD: total variation distance
PMS: Potential/putative methylation signal

## Declarations

## Acknowledgments

We thank Diep Ganguly from Australian National University provides guidance on the use of drought stress data. We thank Professor David Miller from Department of electrical engineering for helpful discussions.

## Funding

The work was supported by funding from the Bill and Melinda Gates Foundation (OPP1088661).

## Availability of data and materials

The source code for all of analysis and visualization, including The Methyl-IT package source code, Network enrichment analysis, R code for all figures are available at the GitLab: https://git.psu.edu/genomath/MethylIT. Seed development methylome dataset, original studied by Kawakatsu et al. (2017) [21], was obtained from the Gene Expression Omnibus (GEO) under accession numbers GSE68132. Drought stress methylome dataset was obtained from the Gene Expression Omnibus (GEO) under accession numbers GSE94075.The differential expressed gene list and express level was obtained Crisp et al. [23].

## Authors’ contributions

RS developed the application of the information thermodynamic theory on cytosine DNA methylation and conducted mathematical and computational biology analyses, XY participated in methylome data analysis, JRB conducted computation, HK conducted the NBEA, NEAT analysis. SM designed experiments, participated in data analysis and wrote manuscript.

## Competing interests

Not applicable

## Consent for publication

Not applicable

## Ethics approval and consent to participate

Not applicable

## Additional files

Additional file 1: Table.S1 DMGs from Arabidopsis seed development dataset.

Additional file 2: Table.S2 List of seed develoment DMGs found_in networks based on NEAT.

Additional file 3: Figure.S1 Interaction network built for the seed development DMGs in networks identified with NEAT using GeneMNIA.

Additional file 4: Table.S3 Enriched network from seed development DMGs (with minimum coverage 10 reads).

Additional file 5: Table.S4 List of 6669 DMGs identifed in the drought stress experiment.

Additional File 6: Table. S5 DMGs in the enriched networks identified by NBEA for the drought stress data.

## References

1. Calarco JP, Borges F, Donoghue MTA, Van Ex F, Jullien PE, Lopes T, Gardner R, Berger F, Feijo JA, Becker JD et al: Reprogramming of DNA Methylation in Pollen Guides Epigenetic Inheritance via Small RNA. Cell 2012, 151(1):194–205.

2. Schmitz RJ, Schultz MD, Lewsey MG, O’Malley RC, Urich MA, Libiger O, Schork NJ, Ecker JR: Transgenerational Epigenetic Instability Is a Source of Novel Methylation Variants. Science 2011, 334(6054):369–373.

3. Becker C, Hagmann J, Muller J, Koenig D, Stegle O, Borgwardt K, Weigel D: Spontaneous epigenetic variation in the Arabidopsis thaliana methylome. Nature 2011, 480(7376):245–U127.

4. Matzke MA, Mosher RA: RNA-directed DNA methylation: an epigenetic pathway of increasing complexity (vol 15, 394, 2014). Nat Rev Genet 2014, 15(8).

5. Crisp PA, Ganguly D, Eichten SR, Borevitz JO, Pogson BJ: Reconsidering plant memory: Intersections between stress recovery, RNA turnover, and epigenetics. Sci Adv 2016, 2(2).

6. Kinoshita T, Seki M: Epigenetic Memory for Stress Response and Adaptation in Plants. Plant Cell Physiol 2014, 55(11):1859–1863.

7. Colaneri AC, Jones AM: Genome-Wide Quantitative Identification of DNA Differentially Methylated Sites in Arabidopsis Seedlings Growing at Different Water Potential. Plos One 2013, 8(4).

8. Severin PM, Zou X, Gaub HE, Schulten K: Cytosine methylation alters DNA mechanical properties. Nucleic Acids Res 2011, 39(20):8740–8751.

9. Osakabe A, Adachi F, Arimura Y, Maehara K, Ohkawa Y, Kurumizaka H: Influence of DNA methylation on positioning and DNA flexibility of nucleosomes with pericentric satellite DNA. Open Biol 2015, 5(10).

10. Yusufaly TI, Li Y, Olson WK: 5-Methylation of cytosine in CG:CG base-pair steps: a physicochemical mechanism for the epigenetic control of DNA nanomechanics. J Phys Chem B 2013, 117(51):16436–16442.

11. Jenkinson G, Pujadas E, Goutsias J, Feinberg AP: Potential energy landscapes identify the information-theoretic nature of the epigenome. Nat Genet 2017, 49(5):719-+.

12. Sanchez R, Mackenzie SA: Information Thermodynamics of Cytosine DNA Methylation. Plos One 2016, 11(3).

13. Sanchez R, Mackenzie SA: Genome-Wide Discriminatory Information Patterns of Cytosine DNA Methylation. Int J Mol Sci 2016, 17(6).

14. Greiner M, Pfeiffer D, Smith RD: Principles and practical application of the receiver-operating characteristic analysis for diagnostic tests. Prev Vet Med 2000, 45(1–2):23–41.

15. Carter JV, Pan J, Rai SN, Galandiuk S: ROC-ing along: Evaluation and interpretation of receiver operating characteristic curves. Surgery 2016, 159(6):1638–1645.

16. Harpaz R, DuMouchel W, LePendu P, Bauer-Mehren A, Ryan P, Shah NH: Performance of pharmacovigilance signal-detection algorithms for the FDA adverse event reporting system. Clin Pharmacol Ther 2013, 93(6):539–546.

17. Kruspe S, Dickey DD, Urak KT, Blanco GN, Miller MJ, Clark KC, Burghardt E, Gutierrez WR, Phadke SD, Kamboj S et al: Rapid and Sensitive Detection of Breast Cancer Cells in Patient Blood with Nuclease-Activated Probe Technology. Mol Ther Nucleic Acids 2017, 8:542–557.

18. Kay SM: Fundamentals of Statistical Signal Processing, Volume II: Detection Theory, 1 edition edn; 1998.

19. Wu H, Xu T, Feng H, Chen L, Li B, Yao B, Qin Z, Jin P, Conneely KN: Detection of differentially methylated regions from whole-genome bisulfite sequencing data without replicates. Nucleic Acids Res 2015, 43(21):e141.

20. Schultz MD, He Y, Whitaker JW, Hariharan M, Mukamel EA, Leung D, Rajagopal N, Nery JR, Urich MA, Chen H et al: Human body epigenome maps reveal noncanonical DNA methylation variation. Nature 2015, 523(7559):212–216.

21. Kawakatsu T, Nery JR, Castanon R, Ecker JR: Dynamic DNA methylation reconfiguration during seed development and germination. Genome Biol 2017, 18(1):171.

22. Schmitz RJ, Schultz MD, Urich MA, Nery JR, Pelizzola M, Libiger O, Alix A, McCosh RB, Chen H, Schork NJ et al: Patterns of population epigenomic diversity. Nature 2013, 495(7440):193–198.

23. Crisp PA, Ganguly DR, Smith AB, Murray KD, Estavillo GM, Searle I, Ford E, Bogdanovic O, Lister R, Borevitz JO et al: Rapid Recovery Gene Downregulation during Excess-Light Stress and Recovery in Arabidopsis. Plant Cell 2017, 29(8):1836–1863.

24. Ganguly DR, Crisp PA, Eichten SR, Pogson BJ: The Arabidopsis DNA Methylome Is Stable under Transgenerational Drought Stress. Plant Physiol 2017, 175(4):1893–1912.

25. Basu A, Mandal A, Pardo L: Hypothesis testing for two discrete populations based on the Hellinger distance. Stat Probabil Lett 2010, 80(3–4):206–214.

26. Vaart A: Asymptotic Statistics: Cambridge University Press; 1998.

27. Mónica López-Ratón MXR-Á, Carmen Cadarso-Suárez, Francisco Gude-Sampedro: OptimalCutpoints: An R Package for Selecting Optimal Cutpoints in Diagnostic Tests. Journal of statistical software 2014, Vol 61 (2014)(Issue 8):4896.

28. Perkins W, Tygert M, Ward R: Computing the confidence levels for a root-mean-square test of goodness-of-fit. Applied Mathematics and Computation 2011, 217(22):9072–9084.

29. Le BH, Cheng C, Bui AQ, Wagmaister JA, Henry KF, Pelletier J, Kwong L, Belmonte M, Kirkbride R, Horvath S et al: Global analysis of gene activity during Arabidopsis seed development and identification of seed-specific transcription factors. Proc Natl Acad Sci U S A 2010, 107(18):8063–8070.

30. Bassel GW, Lan H, Glaab E, Gibbs DJ, Gerjets T, Krasnogor N, Bonner AJ, Holdsworth MJ, Provart NJ: Genome-wide network model capturing seed germination reveals coordinated regulation of plant cellular phase transitions. Proc Natl Acad Sci U S A 2011, 108(23):9709–9714.

31. Mostafavi S, Ray D, Warde-Farley D, Grouios C, Morris Q: GeneMANIA: a real-time multiple association network integration algorithm for predicting gene function. Genome Biol 2008, 9 Suppl 1:S4.

32. Krzywinska E, Bucholc M, Kulik A, Ciesielski A, Lichocka M, Debski J, Ludwikow A, Dadlez M, Rodriguez PL, Dobrowolska G: Phosphatase ABI1 and okadaic acid-sensitive phosphoprotein phosphatases inhibit salt stress-activated SnRK2.4 kinase. Bmc Plant Biol 2016, 16(1):136.

33. Yoshida T, Fujita Y, Maruyama K, Mogami J, Todaka D, Shinozaki K, Yamaguchi-Shinozaki K: Four Arabidopsis AREB/ABF transcription factors function predominantly in gene expression downstream of SnRK2 kinases in abscisic acid signalling in response to osmotic stress. Plant Cell Environ 2015, 38(1):35–49.

34. Yoshida T, Fujita Y, Sayama H, Kidokoro S, Maruyama K, Mizoi J, Shinozaki K, Yamaguchi-Shinozaki K: AREB1, AREB2, and ABF3 are master transcription factors that cooperatively regulate ABRE-dependent ABA signaling involved in drought stress tolerance and require ABA for full activation. Plant J 2010, 61(4):672–685.

35. Yakir E, Hilman D, Hassidim M, Green RM: CIRCADIAN CLOCK ASSOCIATED1 transcript stability and the entrainment of the circadian clock in Arabidopsis. Plant Physiol 2007, 145(3):925–932.

36. Lefebvre A, Mauffret O, el Antri S, Monnot M, Lescot E, Fermandjian S: Sequence dependent effects of CpG cytosine methylation. A joint 1H-NMR and 31P-NMR study. Eur J Biochem 1995, 229(2):445–454.

37. Nathan D, Crothers DM: Bending and flexibility of methylated and unmethylated EcoRI DNA. J Mol Biol 2002, 316(1):7–17.

38. Huang SC, Ecker JR: Piecing together cis-regulatory networks: insights from epigenomics studies in plants. Wiley Interdiscip Rev Syst Biol Med 2017.

39. Krueger F, Andrews SR: Bismark: a flexible aligner and methylation caller for Bisulfite-Seq applications. Bioinformatics 2011, 27(11):1571–1572.

40. Prezza N, Vezzi F, Kaller M, Policriti A: Fast, accurate, and lightweight analysis of BS-treated reads with ERNE 2. Bmc Bioinformatics 2016, 17.

41. Sason I, Verdu S: f-Divergence Inequalities. Ieee Transactions on Information Theory 2016, 62(11):5973–6006.

42. R_Core_Team: A language and environment for statistical computing. 2016.

43. Hippenstiel RD: Detection Theory: Applications and Digital Signal Processing. CRC Press 2001.

44. Stanislaw H, Todorov N: Calculation of signal detection theory measures. Behav Res Meth Ins C 1999, 31(1):137–149.

45. Youden WJ: Index for rating diagnostic tests. Cancer 1950, 3(1):32–35.

46. Perkins NJ, Schisterman EF: The inconsistency of “optimal” cutpoints obtained using two criteria based on the receiver operating characteristic curve. Am J Epidemiol 2006, 163(7):670–675.

47. Carstensen B, Plummer, M., Laara, E. & Hills, M.: Epi:A Package for Statistical Analysis in Epidemiology. R package version 27 2016.

48. Love MI, Huber W, Anders S: Moderated estimation of fold change and dispersion for RNA-seq data with DESeq2. Genome Biol 2014, 15(12).

49. William Perkins MT, Rachel Ward: Computing the confidence levels for a root-mean-square test of goodness-of-fit. Applied Mathematics and Computation 2011, Volume 217(issue 22):Pages 9072–9084.

50. He Y, Gorkin DU, Dickel DE, Nery JR, Castanon RG, Lee AY, Shen Y, Visel A, Pennacchio LA, Ren B et al: Improved regulatory element prediction based on tissue-specific local epigenomic signatures. Proc Natl Acad Sci U S A 2017, 114(9):E1633–E1640.

51. F. Liese IV: On Divergences and Informations in Statistics and Information Theory. IEEE Transactions on Information Theory 2006, Volume: 52(Issue: 10):4394–4412.

52. Geistlinger L, Csaba G, Zimmer R: Bioconductor’s EnrichmentBrowser: seamless navigation through combined results of set- & network-based enrichment analysis. Bmc Bioinformatics 2016, 17.

53. Signorelli M, Vinciotti V, Wit EC: NEAT: an efficient network enrichment analysis test. Bmc Bioinformatics 2016, 17.

